# Dynamics and activation of the *Legionella* Type IV secretion system

**DOI:** 10.64898/2026.10.19.760228

**Authors:** Samira Heydari, Jian Yue, Zi Yang, Sheng Shu, Hongpeng Wang, Wei Mi, Craig Roy, Jun Liu

## Abstract

*Legionella pneumophila* uses the Dot/Icm type IV secretion system (T4SS) to translocate hundreds of effector proteins into host cells, yet how this transenvelope machine assembles and becomes secretion competent remains poorly understood. Here, using cryo-electron tomography and *in situ* single-particle cryo-electron microscopy, we resolve distinct structural and functional states of the native Dot/Icm T4SS. Unexpectedly, the T4SS predominantly occupies a resting state containing the outer-membrane core complex but lacking the inner-membrane complex, including DotA. Recruitment of inner-membrane components establishes a fully assembled but secretion-inactive primed state. A 3.6-Å cryo-EM structure of the DotO–DotI–DotJ–IcmT complex, integrated with *in situ* reconstructions, reveals how membrane-associated components organize DotO into a hexamer-of-dimers ATPase platform. Upon host contact, the T4SS undergoes extensive conformational remodeling while preserving its transenvelope scaffold. Activation involves dilation of the DotG outer-membrane gate and transformation of the DotA–IcmX protochannel into an open translocation conduit that extends beyond the bacterial surface. Substrate-associated state exhibit further channel expansion, revealing structural plasticity associated with effector translocation. During macrophage infection, the extracellular conduit bridges the bacterial and host membranes, establishing a direct pathway for effector delivery. Together, our findings reveal a stepwise mechanism of Dot/Icm T4SS channel assembly and activation, in which coordinated remodeling of the secretion channel converts a resting apparatus into a host-engaged translocation machine.

## Introduction

*Legionella pneumophila* is an intracellular Gram-negative bacterial pathogen that causes a severe form of pneumonia known as Legionnaires’ disease ^1^. Following uptake by host phagocytic cells, *L. pneumophila* establishes a replication-permissive niche within a specialized compartment, the *Legionella*-containing vacuole (LCV), by extensively reprogramming host membrane trafficking and immune signaling pathways ^2,3^. This remodeling is mediated by the Dot/Icm type IV secretion system (T4SS), a large transenvelope nanomachine that translocates hundreds of functionally diverse effector proteins into host cells ^4,5^. Dot/Icm-mediated effector translocation is essential for bacterial intracellular survival, replication, and virulence ^6^. Composed of at least 27 different proteins, the Dot/Icm T4SS spans the inner membrane, periplasm, and outer membrane, making it one of the most architecturally complex bacterial secretion systems described to date ^7,8^. Previous structural studies have revealed a modular organization consisting of an outer membrane core complex (OMCC) and inner membrane complex (IMC), which together form the envelope-spanning secretion apparatus ^9–12^.

Cryo-electron tomography (cryo-ET) studies visualized successive stages of Dot/Icm T4SS biogenesis within intact bacterial cells, revealing a hierarchical assembly pathway involving the sequential incorporation of distinct subassemblies ^9^. Notably, the OMCC and IMC required separate refinement to resolve their structures, suggesting substantial conformational flexibility within the intact secretion machinery in addition to compositional heterogeneity associated with assembly intermediates. More recently, we identified inactive and active conformations characterized by closed and open secretion channels, respectively ^13^. The closed state was resolved using a DotB^E191K^ mutant strain ^14^, in which ATPase-deficient DotB ^15^ stably associates with DotO, enriching fully assembled Dot/Icm complexes. This stabilization enabled identification of the previously unassigned channel components DotA and IcmX ^13^. However, because the DotB^E191K^ mutant is defective in ATP hydrolysis and secretion, it preferentially stabilizes the fully assembled inactive state and cannot capture the full conformational landscape of the wild-type secretion system. Consequently, how the native Dot/Icm T4SS transitions from assembly intermediates to secretion-competent states, and how these transitions are coordinated with substrate translocation, remain poorly understood.

Here, we investigate the structural dynamics and activation of the native Dot/Icm T4SS in wild-type *L. pneumophila* by integrating cryo-focused ion beam (cryo-FIB) milling and cryo-ET imaging with *in situ* single-particle cryo-EM. We identify distinct resting, primed, and active states, revealing structural transitions associated with secretion-system maturation and activation. A high-resolution cryo-EM structure of the DotO–DotI–DotJ–IcmT complex, integrated with *in situ* cryo-ET maps, defines the molecular architecture of the IMC and completes the structural model of the primed Dot/Icm T4SS. Furthermore, *in situ* structures reveal extensive remodeling of the central secretion apparatus during activation, including channel opening and formation of an extracellular conduit extending beyond the bacterial surface. By combining cryo-FIB milling with cryo-ET imaging of *L. pneumophila*-infected macrophages, we directly visualize the engagement of the activated Dot/Icm T4SS with the host membrane in its native infection context. Collectively, our findings reveal the structural transitions underlying Dot/Icm T4SS maturation and activation and establish how extensive remodeling of the secretion apparatus enables the formation of a translocation conduit at the host–pathogen interface. More broadly, this work provides mechanistic insight into how a dynamic bacterial secretion nanomachine coordinates structural remodeling with effector translocation during intracellular infection.

## Results

### Dot/Icm T4SS predominantly adopts a DotA-deficient resting state in wild-type *L. pneumophila*

To determine the native structural organization and conformational variability of the Dot/Icm T4SS in wild-type *L. pneumophila*, we visualized intact bacterial cells by cryo-ET (Fig. 1a). A total of 403 tilt series were collected, from which 1,499 Dot/Icm T4SS particles were identified and subjected to subtomogram averaging and classification. The predominant population (970 particles; 64.7%) lacked detectable DotA density but retained the disk, dome, periplasmic ring, and IcmX (Fig. 1b). We designated this population the resting state. The remaining particles exhibited detectable DotA density and were classified as the primed state. Further classification of the DotA-containing particles revealed two distinct conformations with similar overall architectures but markedly different axial dimensions. One exhibited an outer-to-inner membrane spacing of approximately 30 nm, whereas the other displayed an increased spacing of approximately 34 nm (Fig. 1c,d). We designated these as relaxed and expanded conformations, respectively. Notably, the expanded conformation was accompanied by pronounced inward curvature of the inner membrane, suggesting that axial expansion of the secretion apparatus is coupled to localized remodeling of the bacterial envelope. Together, these observations reveal substantial structural heterogeneity within the native Dot/Icm T4SS and suggest that DotA incorporation into the central channel is associated with conformational changes that may facilitate recruitment of the cytoplasmic ATPase machinery and prime the secretion apparatus for subsequent activation.

**Fig. 1.**
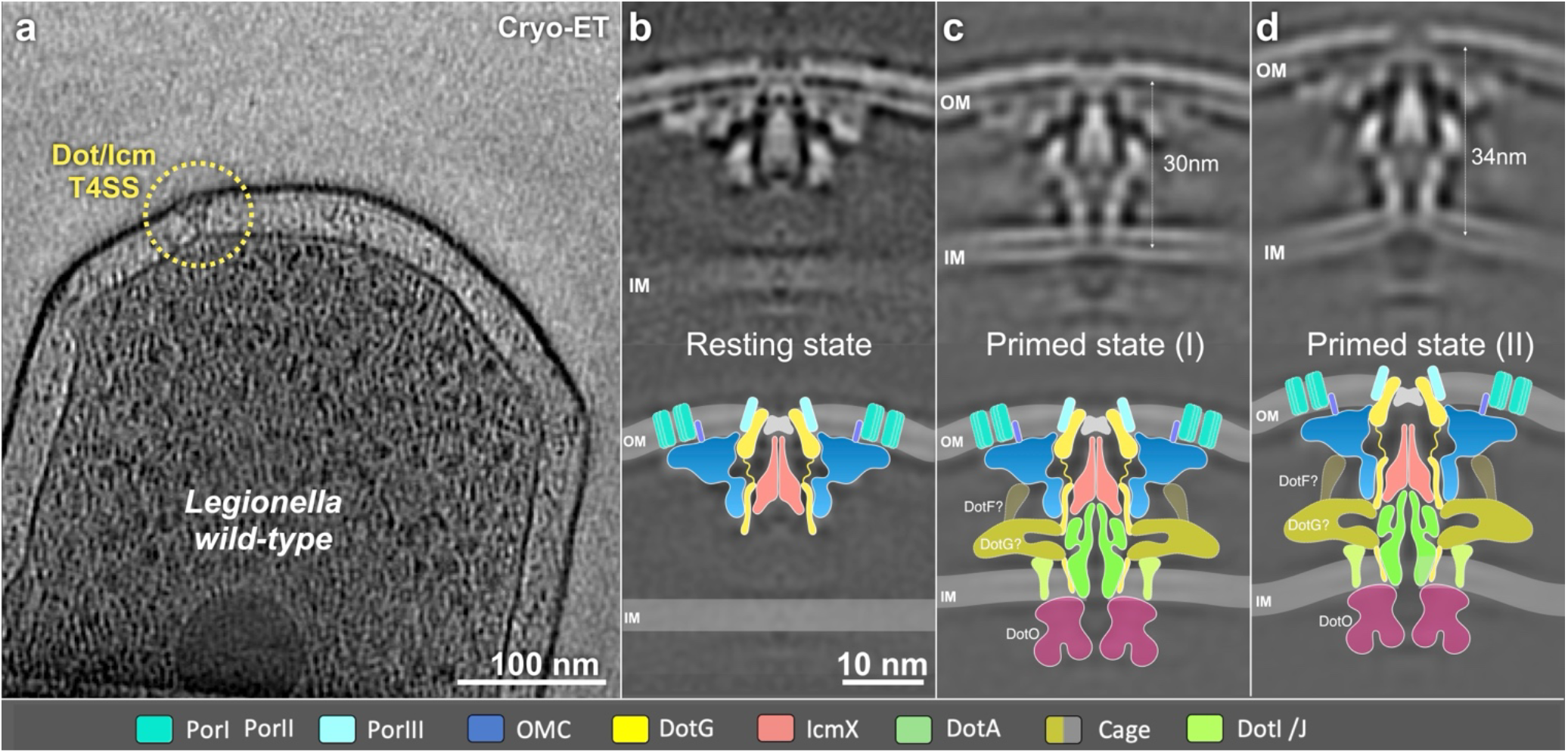
Cryo-ET reveal the DotA-deficient state as the predominant Dot/Icm T4SS architecture in wild-type *L. pneumophila*. **a,** Representative tomographic slice of a wild-type (WT) cell showing a polar Dot/Icm T4SS (dashed circle). **b–d,** Subtomogram averages and corresponding models of three distinct Dot/Icm T4SS populations. The predominant population (64.7%) lacks resolved DotA density (b), whereas two less abundant populations exhibit well-resolved DotA density and distinct conformations characterized by outer membrane–inner membrane (OM–IM) spacings of ∼30 nm (c) and ∼34 nm (d), respectively.

### Molecular architecture of the primed Dot/Icm T4SS

Although *in situ* cryo-EM resolved the DotA-containing central channel of the primed Dot/Icm T4SS ^13^, much of the IMC remained poorly resolved, likely owing to its conformational flexibility. To define the molecular architecture of the IMC, we purified the native DotO-associated complex for single-particle cryo-EM analysis. A C-terminal FLAG affinity tag was introduced at the chromosomal dotO locus, and detergent solubilization followed by affinity purification yielded a stable complex comprising DotO, DotI, DotJ, and IcmT (Fig. 2a; Extended Data Fig. 1). Cryo-EM reconstruction at approximately 3.6 Å resolution revealed a DotO:DotI:DotJ:IcmT stoichiometry of 2:3:1:1 (Fig. 2c–f; Extended Data Fig. 1). Six copies of this complex could be fitted into the corresponding cytoplasmic density of the *in situ* cryo-ET map ^13^, enabling reconstruction of the complete IMC assembly (Fig. 2g,h). In this architecture, six inner DotO subunits form a central hexamer, while six outer DotO subunits extend radially toward the surrounding membrane-associated components. Together, they form a hexamer-of-dimers ATPase platform anchored around the DotA channel in the inner membrane through interactions with DotI, DotJ, and IcmT.

**Fig. 2.**
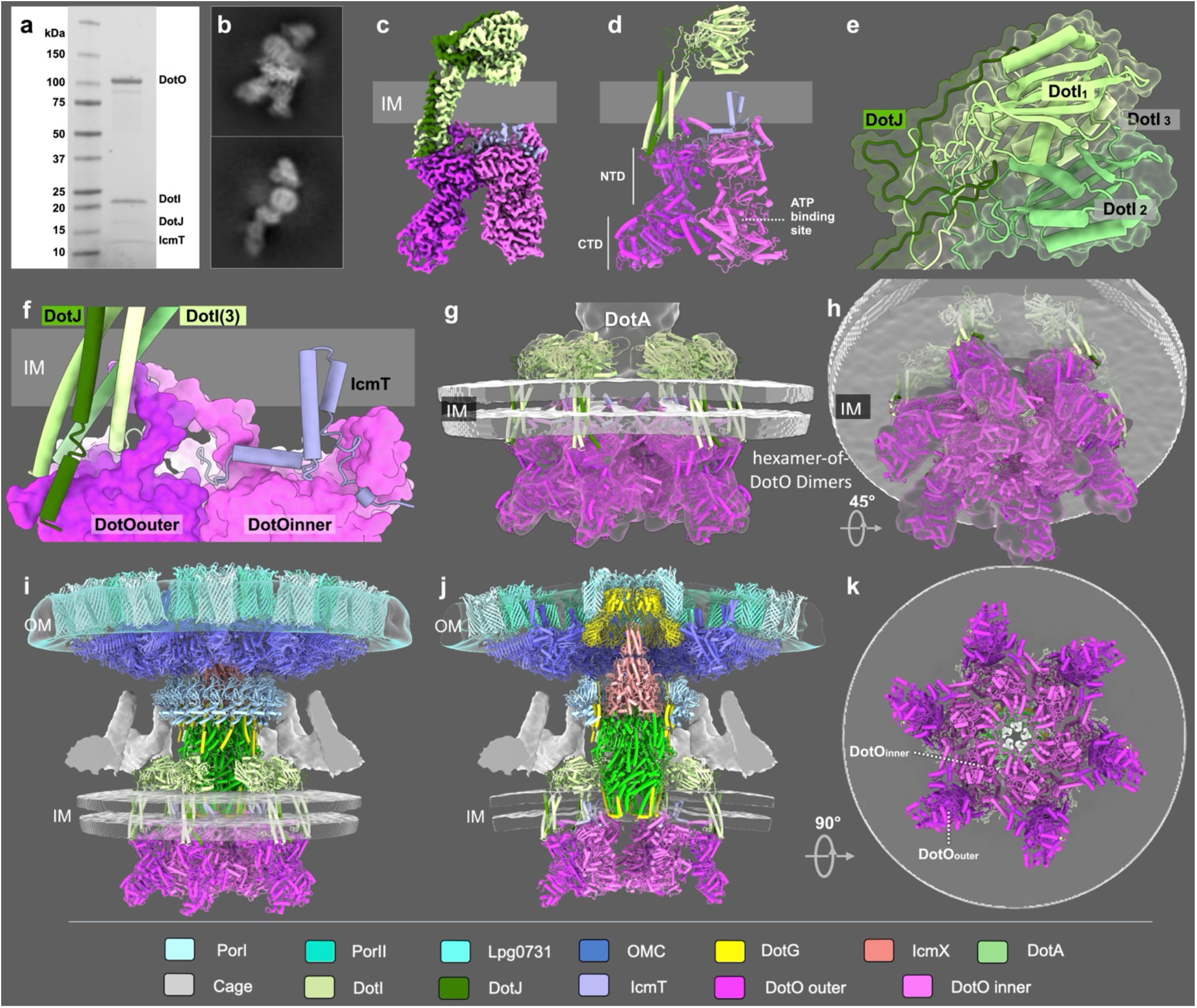
Cryo-EM structure of the inner membrane complex completes the near-atomic architecture of the inactive primed Dot/Icm T4SS. **a,** SDS–PAGE analysis of the affinity-purified inner membrane complex (IMC) containing DotO, DotI, DotJ and IcmT. **b,** Representative 2D class averages of the purified complex. **c,** Cryo-EM density map of the DotO–DotI–DotJ–IcmT complex resolved at ∼3.6 Å, revealing a stoichiometry of 2:3:1:1. **d,** Atomic model highlighting the N-terminal (NTD) and C-terminal (CTD) domains of DotO and the predicted ATP-binding site. **e,** Structural organization of the DotI trimer and its associated DotJ subunit. **f,** Close-up of the membrane-associated interface showing interactions of DotI–DotJ and IcmT with the outer and inner DotO subunits, respectively. **g,h** six copies of the purified IMC fitted into the cryo-ET density to reconstruct the complete IMC assembly. **i, j,** Composite models of the inactive Dot/Icm T4SS integrating the IMC with the envelope-spanning apparatus. k, Bottom view of the DotO hexamer-of-dimers assembly, showing the central inner-DotO hexamer and radially arranged outer DotO subunits.

Within each dimer, the two DotO subunits interact primarily through their N-terminal regions (residues 1–450). The outer DotO subunit is extensively coupled to the DotI–DotJ membrane-associated assembly. Three DotI molecules form a trimeric module whose periplasmic domains contribute to an arch-like structure above the inner membrane, while their transmembrane and cytoplasmic regions associate with DotJ and extend toward the membrane-proximal surface of DotO (Fig. 2e,f). In contrast, IcmT primarily contacts the inner DotO subunit and is positioned adjacent to the DotA pentamer, providing an additional membrane-proximal connection between the ATPase platform and the central secretion apparatus (Fig. 2f). Thus, DotI–DotJ and IcmT preferentially engage the outer and inner DotO subunits, respectively, establishing an asymmetric DotO dimer that serves as the building block of the IMC.

DotO is the major VirB4-family ATPase of the Dot/Icm T4SS and is positioned to couple nucleotide turnover to conformational changes within the secretion machinery ^16,17^. The predicted nucleotide-binding pocket lies within the central region of each DotO subunit, adjacent to the C-terminal region (residues 916–1009) (Fig. 2d; Extended Data Fig. 2,3). Notably, the C-terminal regions of neighboring DotO molecules are positioned in close proximity within the oligomeric assembly, suggesting potential coupling between adjacent catalytic regions. This organization establishes a structural connection between the catalytic regions of DotO and its membrane-associated N-terminal scaffold, providing a potential route through which ATP-dependent conformational changes could be transmitted to the DotA-containing secretion channel.

In addition to resolving the IMC, we identified Lpg0731 as a component of Porin III, an outer-membrane ring surrounding the DotG dome. Deletion of lpg0731 resulted in the loss of Porin III density, supporting the assignment of Lpg0731 to this ring (Extended Data Fig. 4). Together, the identification of Lpg0731, the high-resolution structure of the DotO–DotI–DotJ–IcmT complex, and our previously determined near-atomic *in situ* architecture of the periplasmic apparatus ^13^ enable reconstruction of an integrated structural model of the primed Dot/Icm T4SS, extending from the IMC to the OMCC (Fig. 2i–k; Extended Data Movie S1). This model defines the molecular organization of the Dot/Icm T4SS in primed state and provides a structural foundation for understanding the conformational changes associated with Dot/Icm T4SS activation.

### Dot/Icm T4SS population remains predominantly in the resting state upon secretion stimulation

To capture the active Dot/Icm T4SS, we stimulated secretion by incubating *L. pneumophila* with macrophage lysate ^13^ before vitrification (Fig. 3a) and collected 76,493 micrographs with *in situ* cryo-EM (Fig. 3b,c). 65,146 Dot/Icm T4SS particles were identified through combined manual and Topaz-assisted particle picking (Extended Data Fig. 5). The consensus reconstruction, refined with C13 symmetry, lacked resolved DotA density, resembling the predominant resting state observed in unstimulated cells.

**Fig. 3.**
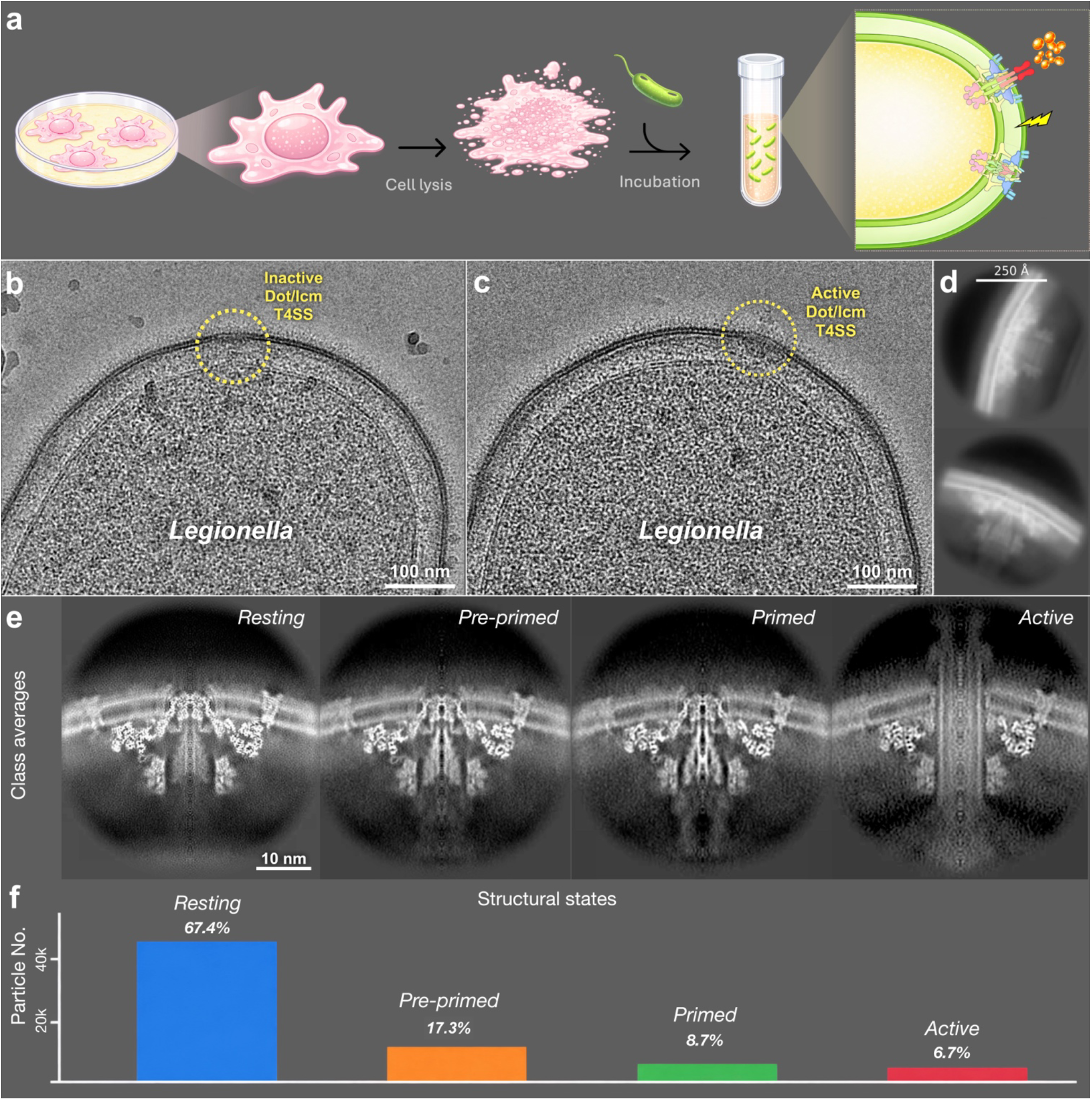
*In situ* cryo-EM reveals distinct structural states of the *L. pneumophila* Dot/Icm T4SS following exposure to host-cell lysate. **a,** Schematic illustration of the sample preparation workflow for stimulating secretion, including host-cell lysis and subsequent incubation with *L. pneumophila*. **b,c,** Representative cryo-EM micrographs showing inactive (b) and active (c) Dot/Icm T4SS assemblies, highlighted by yellow dashed circles. **d,** Representative 2D class averages of the Dot/Icm T4SS. **e,f,** Representative class averages and corresponding particle distributions of four distinct structural states: resting (67.4%), pre-primed (17.3%), primed (8.7%) and active (6.7%).

To distinguish conformational states of the central secretion apparatus, we subtracted the densities corresponding to the OMCC, PR, and dome, followed by focused 3D classification of C13 symmetry-expanded particles using a mask encompassing IcmX and DotA. This analysis identified four distinct structural states that differed primarily in the organization of the central secretion pathway (Fig. 3e; Extended Data Fig. 5). Particles assigned to each class were subsequently re-extracted and reconstructed without signal subtraction to visualize the intact secretion complexes.

The predominant population (43,902 particles; 67.4%) lacked detectable DotA density while retaining the disk, dome, PR, and IcmX, consistent with the resting state. A second population (11,239 particles; 17.3%) exhibited weak, axially restricted density at the expected position of DotA that could not be further resolved, suggesting substantial conformational heterogeneity. We designated this population the pre-primed state, in which DotA may associate with the secretion apparatus but remains conformationally mobile. A third population (5,640 particles; 8.7%) displayed well-resolved DotA density, defining the primed state. Notably, despite containing approximately half as many particles as the pre-primed population, this class yielded substantially better-resolved DotA density, indicating that the improved structural order reflects a distinct conformational state rather than simply differences in particle number. The remaining population (4,365 particles; 6.7%) represented the active state, characterized by extensive remodeling of the central secretion apparatus and formation of an extracellular extension projecting beyond the bacterial surface, as described below.

Thus, even following secretion stimulation, approximately two-thirds of Dot/Icm T4SS complexes remain in the resting state, whereas progressively smaller populations occupy the pre-primed, primed, and active states. These findings suggest that DotA is not a constitutively ordered component of the secretion apparatus but instead undergoes progressive stabilization during maturation of the central secretion channel. The transition from a poorly ordered to a well-resolved DotA-containing channel may facilitate its functional coupling to the cytoplasmic ATPase machinery, thereby preparing the complex for secretion. Notably, the low abundance of primed and active complexes suggests that formation and stabilization of the DotA-containing protochannel may represent a key regulatory step preceding full activation. Together, these observations reveal a heterogeneous population of native Dot/Icm T4SS complexes and support a model in which progressive organization of the central secretion apparatus precedes activation and effector translocation.

### The Dot/Icm T4SS undergoes extensive channel remodeling and forms a variable-length extracellular conduit upon activation

The C13-symmetric disk and C18-symmetric PR of the active Dot/Icm T4SS were reconstructed at resolutions of 4.2 Å and 6.94 Å, respectively (Extended Data Fig. 5). Comparison of the primed and active structures revealed that the overall architectures of the OMCC disk and PR remain largely preserved upon activation, whereas extensive structural remodeling occurs along the central secretion axis. In the active state, the secretion channel forms a continuous conduit extending from the inner membrane through the periplasmic apparatus and outer membrane into the extracellular space (Fig. 4a, b; Movie S2). Remarkably, the extracellular extension terminates in a distinctive dumbbell-shaped tip, suggesting a potential role in host-membrane engagement during effector translocation, although its molecular composition and precise function remain to be determined.

**Fig. 4.**
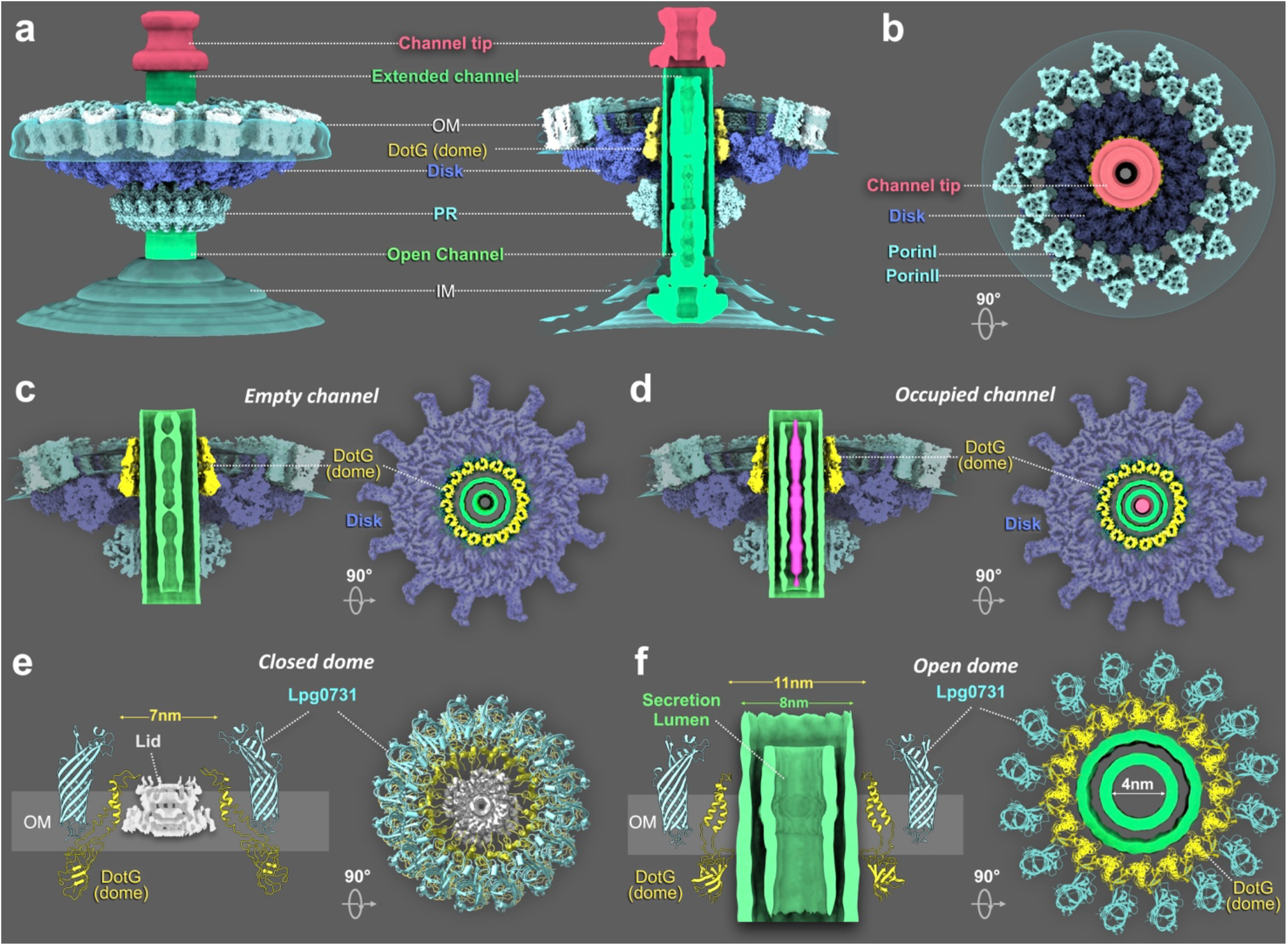
Structural characterization of the active Dot/Icm T4SS reveals an open extracellular secretion conduit. **a**, Surface-rendered views of the active Dot/Icm T4SS reconstruction illustrating the overall architecture of the secretion apparatus, including the open secretion channel, its extracellular extension, and the distal channel tip. **b**, Top view of the active Dot/Icm T4SS structure highlighting the open secretion channel within the OMCC disk and the surrounding Porin I and Porin II assemblies. **c**,**d**, Two distinct active Dot/Icm T4SS classes revealing an empty (c) or occupied (d) secretion conduit. **e**,**f** Structural comparison illustrating activation-dependent remodeling of the outer membrane (OM) secretion gate. In the inactive state (e), the constricted DotG dome is occluded by a central lid and surrounded by an ordered Porin III/Lpg0731 assembly, forming a closed distal gate. Upon activation (f), the DotG dome aperture expands from ∼7 to ∼11 nm, accompanied by loss of the central lid and increased disorder of the surrounding Porin III/Lpg0731 assembly. Together, these structural changes open the OM secretion gate and establish a continuous extracellular conduit for substrate translocation.

To investigate whether the active secretion channel undergoes further conformational changes during substrate translocation, we performed focused 3D classification using a mask encompassing the central conduit. This analysis revealed two distinct channel architectures: one with a continuous open lumen lacking detectable internal density and another containing an elongated density within the central lumen (Fig. 4c,d; Extended Data Fig. 6). Mapping particles from each class back to the original micrographs revealed additional density immediately above the extracellular extension in particles assigned to the empty-lumen class (Extended Data Fig. 6). This density may represent substrate that has exited the secretion channel, suggesting that the empty-lumen and lumen-occupied classes correspond to post-translocation and substrate-engaged states, respectively. Notably, the elongated density extends along the longitudinal axis of the channel, consistent with the possibility that effector proteins adopt an extended conformation during translocation.

Comparison of these two channel architectures revealed localized dilation associated with substrate occupancy. In the empty-lumen state, the inner wall defines a translocation pathway approximately 3 nm in diameter. By contrast, the lumen expands to approximately 4 nm in the substrate-engaged state, while the outer diameter of the conduit remains largely unchanged at approximately 8 nm (Fig. 4c,d). These findings reveal substantial conformational plasticity of the inner channel wall within a comparatively stable outer scaffold, suggesting that localized dilation enables the secretion conduit to accommodate diverse substrates while maintaining its overall structural integrity.

Focused classification of the extracellular portion of the complex further revealed variation in the conduit length (15–20 nm), while the distal tip complex remained largely invariant. Rare particles with substantially longer extensions were also observed, suggesting that the extracellular conduit may undergo further elongation (Extended Data Fig. 6). Together, these observations indicate that the activated secretion conduit exhibits structural flexibility in both its lumen diameter and extracellular length.

Activation was also accompanied by pronounced remodeling of the DotG dome and the surrounding Porin III/Lpg0731 ring. In the primed state, the C-terminal helices of DotG converge toward the central axis, defining an approximately 7-nm aperture occluded by a small, unassigned density termed the lid. The Porin III/Lpg0731 ring surrounds the DotG dome, while the lid occupies its central opening. Upon activation, the C-terminal helices of DotG move outward, expanding the aperture from approximately 7 to 11 nm, accompanied by disappearance of the lid density and relief of the central occlusion. Although the Porin III/Lpg0731 ring remains associated with the DotG dome, it becomes substantially less ordered in the active state. These coordinated structural changes suggest a mechanism of outer-membrane gate opening in which DotG dome dilation, lid displacement, and increased conformational disorder of Porin III/Lpg0731 collectively convert the closed primed architecture into an open secretion conduit (Fig. 4e,f).

### The Dot/Icm T4SS builds dynamic, variable-length bridges across the bacterium–host interface

To determine how the activated Dot/Icm T4SS engages the host membrane during infection, we visualized individual secretion complexes within *L. pneumophila*-infected macrophages using cryo-FIB milling and cryo-ET imaging. At sites of close bacterium–host association, activated Dot/Icm T4SS complexes formed extracellular bridges spanning the space between the bacterial outer membrane and the host membrane, establishing a direct physical connection across the host–pathogen interface (Fig. 5; Movie S3). Strikingly, the length of these bridges varied with the local geometry of the host membrane. Longer bridges were associated with relatively flat host membranes, whereas shorter bridges coincided with pronounced host membrane curvature toward the bacterial surface (Fig. 5c–j). These observations reveal a close relationship between the architecture of the extended secretion apparatus and local host-membrane deformation, suggesting that Dot/Icm T4SS engagement is associated with remodeling of the host–pathogen interface. The variable bridge lengths and associated membrane geometries may represent distinct stages of host-membrane engagement. We propose a model in which the activated Dot/Icm T4SS initially establishes contact with the host membrane through an extended extracellular conduit, followed by progressive shortening of the bacterium–host separation and increased curvature of the host membrane toward the bacterial surface. Such coordinated remodeling could transform an extended secretion bridge into a close-contact interface that facilitates efficient effector translocation. Together, these *in situ* observations provide structural evidence that the activated Dot/Icm T4SS forms a physical bridge between bacterial and host membranes and suggests a mechanism by which dynamic remodeling of the secretion conduit and host membrane may facilitate effector delivery during infection.

**Fig. 5.**
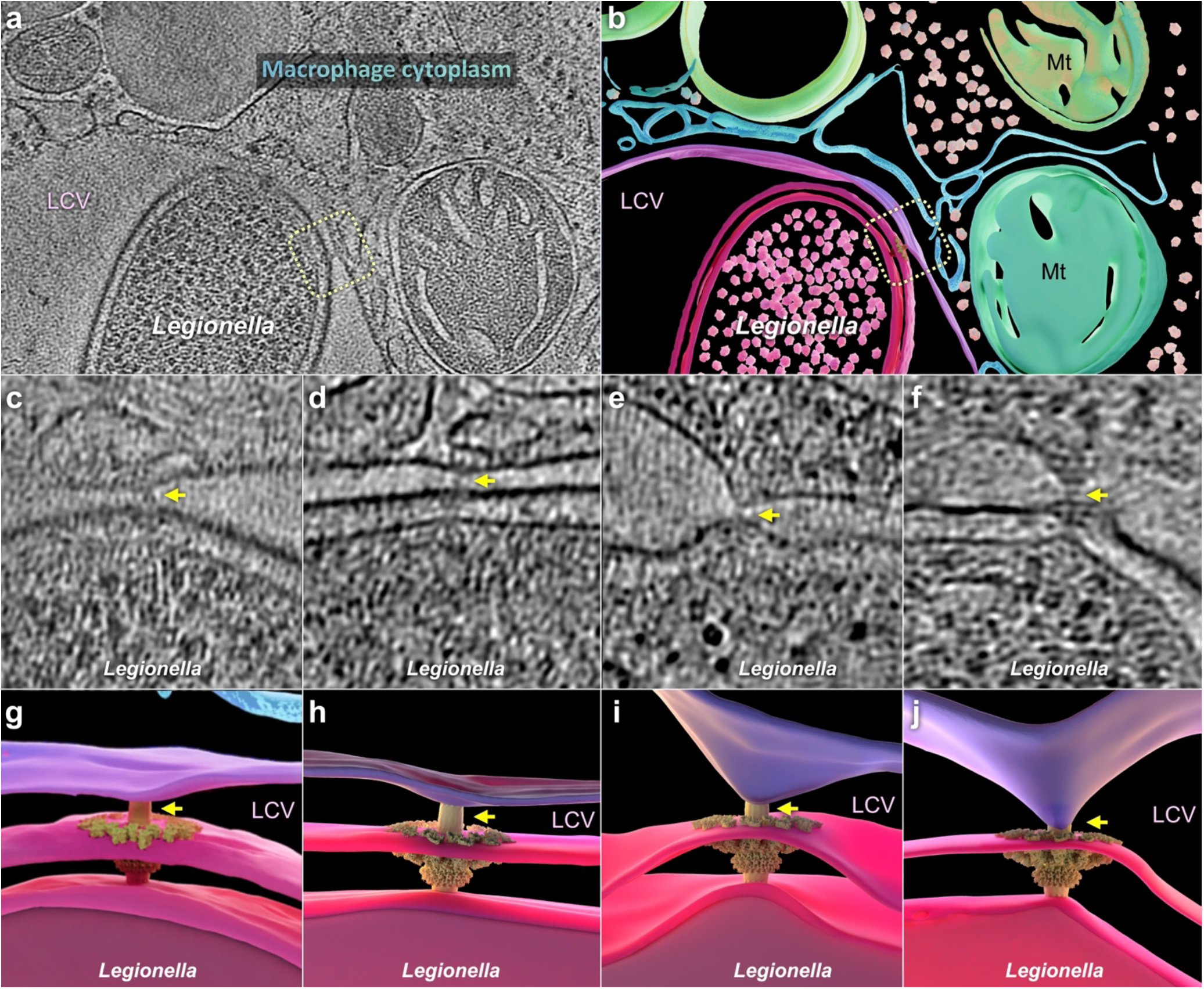
Cryo-FIB/ET captures dynamic bridging of the bacterium–host interface by the Dot/Icm T4SS. **a**, Representative tomographic slice of an *L. pneumophila*-infected macrophage showing an active Dot/Icm T4SS contacting the LCV membrane (dashed box). **b**, 3D segmentation of the tomogram showing the active Dot/Icm T4SS positioned at the bacterium–host membrane interface. **c–f**, Representative cryo-ET tomographic slices illustrating variations in the spacing between the bacterial and LCV membranes while maintaining direct contact (yellow arrows) between the secretion apparatus and the host membrane. **g–j**, Corresponding 3D segmentations of the host-engaged Dot/Icm T4SS assemblies, highlighting variations in bacterium–host membrane spacing and the structural adaptability of the secretion apparatus at the host interface.

## Discussion

T4SSs are widespread bacterial nanomachines and among the most functionally versatile secretion systems, mediating the translocation of proteins and nucleic acids in processes ranging from conjugation to host–pathogen interactions ^18–21^. The Dot/Icm T4SS is particularly elaborate, requiring the coordinated assembly of at least 27 proteins ^4,5,13,22^. Despite substantial advances in defining the architecture of these nanomachines, how such large transenvelope systems transition between functional states to engage and translocate substrates remains a central unsolved question. Here, by integrating *in situ* cryo-EM, cryo-ET and single-particle cryo-EM, we define distinct structural states of the Dot/Icm T4SS in wild-type *L. pneumophila* and reveal progressive remodeling of the secretion pathway during assembly, activation, and substrate translocation.

Our analyses identify the DotA-deficient architecture as the predominant structural population in wild-type *L. pneumophila*, suggesting a physiologically favored resting state from which the machinery progresses toward complete assembly and activation. Together with previous evidence that DotA is incorporated late in the hierarchical assembly pathway ^9^, this finding suggests that stable DotA incorporation represents an important structural transition in T4SS maturation, enabling formation of the fully assembled but inactive primed state. The primed apparatus itself is conformationally dynamic, adopting relaxed and expanded architectures that differ substantially in their trans-envelope dimensions. Notably, the expanded conformation exhibits inner-membrane curvature resembling that of the active apparatus, suggesting that it may represent a pre-activation intermediate. This interpretation is consistent with the association between complete ATPase organization and inner-membrane curvature reported by Dutka and colleagues ^23^. Thus, maturation of the Dot/Icm T4SS appears to involve not simply sequential incorporation of components, but progressive conformational remodeling that prepares the apparatus for activation.

Our structures further define the molecular organization of the IMC within the primed apparatus, providing a high-resolution basis for the hexamer-of-dimers architecture previously observed by cryo-ET ^14,24^. The two DotO subunits within each dimer occupy distinct membrane-associated environments: IcmT primarily engages the inner DotO subunit, whereas the DotI–DotJ module associates with the outer subunit. This asymmetric organization shows an intriguing architectural parallel with the R388 conjugative T4SS, in which the VirB4 homologue TrwK also forms central–peripheral dimers, with VirB3 associated with central TrwK and VirB8 with the peripheral copy ^25^. DotI is structurally related to VirB8, whereas IcmT lacks detectable sequence homology to VirB3 despite occupying an analogous position within the ATPase assembly. Thus, although individual components and interactions have diverged, differential membrane coupling of the two VirB4-family ATPase subunits appears to be preserved across evolutionarily distinct T4SSs. In R388, peripheral TrwK subunits have been proposed to play a predominantly structural role in supporting membrane-associated VirB8 components, whereas the central subunits constitute the central ATPase ^26^. Our structure therefore provides a molecular explanation for the asymmetric DotO dimer and suggests that the hexamer-of-dimers organization represents a conserved architectural framework capable of accommodating distinct membrane-coupling modules across T4SS families.

The DotO architecture also provides a basis for considering how ATP turnover could be coupled to conformational changes within the secretion machinery. The catalytic regions of neighboring DotO subunits are positioned in close proximity within the oligomeric ATPase layer, potentially enabling nucleotide-dependent intersubunit coupling, whereas their N-terminal regions form a more stable scaffold connecting DotO to the membrane-associated DotI–DotJ–IcmT assembly. This organization provides a potential structural route through which ATP-dependent conformational changes could be transmitted from the cytoplasmic ATPase platform toward the membrane-embedded secretion machinery. Determining the nucleotide-dependent conformational cycle of DotO will be important for establishing how ATP hydrolysis drives activation and substrate translocation.

A longstanding question has been how the Dot/Icm T4SS reorganizes to support effector translocation. Our previous *in situ* study captured an active state and showed that activation is accompanied by remodeling of the central secretion axis and formation of a continuous conduit extending beyond the bacterial surface ^13^. The substantially larger *in situ* cryo-EM dataset analyzed here resolves this transition in greater detail. Despite extensive remodeling of the central pathway, the surrounding OMCC disk and periplasmic ring remain largely unchanged, indicating that activation occurs within a comparatively stable envelope-spanning scaffold. By contrast, the central secretion axis undergoes pronounced structural changes that open the outer-membrane gate and generate an extracellular conduit.

A major component of this transition is remodeling of the DotG dome. Previous studies localized the C-terminal VirB10-like domain of DotG to the dome at the distal end of the apparatus ^13,27,28^. In the primed state, the DotG helices converge around an approximately 7-nm aperture occluded by an unassigned lid density and surrounded by the Porin III ring, which we identify as containing Lpg0731. Upon activation, the DotG helices move outward to expand the aperture to approximately 11 nm, the lid density disappears, and the Porin III/Lpg0731 ring becomes substantially less ordered. These coordinated changes convert the closed distal architecture into an open gate through which the central conduit extends beyond the bacterial surface. This transition is consistent with the proposed role of VirB10-family proteins as trans-envelope sensors linking cytoplasmic ATPase activity to conformational changes at the outer membrane ^29^. The closed and open DotG architectures therefore provide a structural basis for regulated outer-membrane gating, although how ATPase activity is energetically coupled to this transition and what role Lpg0731 plays in gate regulation remains unresolved.

The active conduit itself exhibits additional structural plasticity during substrate translocation. Focused classification resolved lumen-empty and lumen-occupied populations, with the latter containing an elongated density along the central axis that may represent substrate captured during passage through the secretion channel. Strikingly, lumen occupancy is accompanied by expansion of the inner channel from approximately 3 to 4 nm, whereas the ∼8-nm outer wall remains largely unchanged. This organization suggests that the conduit combines a comparatively rigid outer shell with a deformable inner lining, allowing localized dilation of the translocation pathway without compromising overall channel integrity. Such adaptability may be particularly important for the Dot/Icm system, which must accommodate hundreds of structurally diverse effector proteins.

The extracellular portion of the active conduit is also highly dynamic. Focused classification revealed extensions approximately 15–20 nm in length with a nearly invariant distal tip, indicating that length variation occurs primarily within the connecting channel. Rare particles with substantially longer extensions suggest an even greater capacity for elongation. This variability recalls extracellular appendages of conjugative T4SSs and the *Helicobacter pylori* Cag T4SS ^30,31^. However, the continuity of the Dot/Icm extension with the central secretion conduit and preservation of a distinct terminal tip suggests that it represents a specialized extension of the translocation pathway rather than a conventional T4SS pilus.

Our cryo-FIB/ET analysis places this structural variability in the context of host engagement during intracellular infection. Longer Dot/Icm bridges contact relatively flat regions of the host membrane, whereas shorter bridges are associated with pronounced membrane curvature toward the bacterial surface. These structures may represent different stages of host engagement, in which an extended conduit establishes initial contact and subsequent shortening of the bacterium–host interface is accompanied by closer membrane apposition. Importantly, these static snapshots do not establish the temporal order or direction of these transitions. Nevertheless, they support a model in which an adjustable extracellular conduit accommodates changes in bacterium–host membrane separation while maintaining contact through a conserved distal tip. Such structural adaptability could allow the Dot/Icm T4SS to maintain a continuous translocation pathway across a dynamically changing bacterium–host interface.

Together, our findings support a model of progressive Dot/Icm T4SS channel remodeling during assembly, activation, substrate translocation and host engagement (Fig. 6). Stable incorporation of DotA and assembly of the cytoplasmic complex establish the primed apparatus. Activation then remodels the central secretion axis, opens the DotG outer-membrane gate and generates an extended extracellular conduit while leaving the surrounding scaffold largely intact. The conduit can adjust its length during host engagement and locally dilate its lumen during substrate passage, providing structural flexibility at both the extracellular and translocation interfaces. These findings reveal the Dot/Icm T4SS not as a static transenvelope machine, but as a dynamic secretion apparatus in which a stable scaffold supports extensive remodeling of the central pathway. Defining the molecular composition of the open channel and distal tip and establishing the temporal and energetic coupling among these transitions, will be important for understanding how the Dot/Icm T4SS delivers its exceptionally diverse effector repertoire across the bacterium–host interface.

**Fig. 6.**
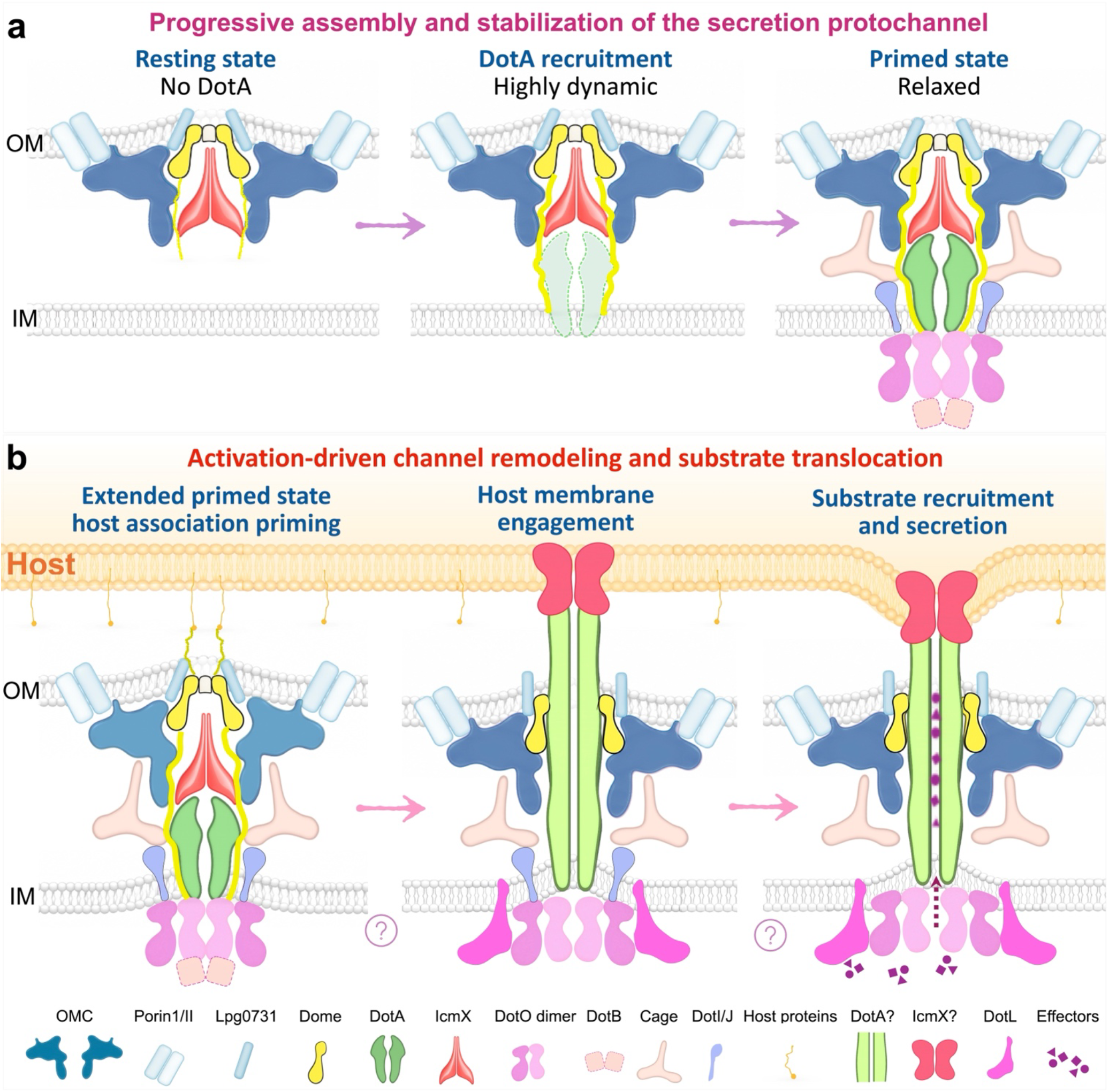
Proposed model for dynamic remodeling of the Dot/Icm T4SS from channel maturation to substrate translocation. **a**, Assembly model illustrating the distinct structural states identified in wild-type *L. pneumophila*. The predominant resting state lacks resolved DotA density. Initial recruitment of DotA is associated with a dynamic configuration, followed by DotA stabilization and assembly of the IMC to establish the primed state. The primed apparatus exhibits conformational plasticity between relaxed and expanded states, characterized by axial extension of the secretion machinery and increased inward curvature of the inner membrane. **b**, Activation model illustrating the proposed progression from host engagement to substrate translocation. Host contact is proposed to trigger extensive remodeling of the central secretion axis, including opening of the outer membrane (OM) gate and formation of a continuous secretion conduit extending beyond the bacterial surface. The extracellular conduit engages the host membrane, establishing a direct bacterium–host connection for substrate translocation. Following initial engagement, the host membrane is proposed to move toward the bacterial OM, reducing the intermembrane distance and potentially facilitating substrate delivery. During active secretion, the conduit may undergo further remodeling, including localized expansion of its lumen to accommodate substrate passage. Arrows indicate the proposed transitions between structural and functional states, whereas question marks denote unresolved molecular components or mechanistic steps.

## Materials and Methods

### Bacterial strains and culture conditions

All experiments were performed using the *L. pneumophila* Philadelphia-1 derivative LP01 wild-type strain. Bacteria were cultured on charcoal yeast extract (CYE) agar plates at 37 °C. Fresh colonies were resuspended in phosphate-buffered saline (PBS) to an optical density (OD600) of approximately 1 immediately prior to cryo-EM grid preparation.

### Preparation of frozen-hydrated specimens

Freshly prepared wild-type *L. pneumophila* suspensions (OD600≈1) were applied to glow-discharged Quantifoil Cu R2/1 300-mesh grids. After incubation for 1 min, grids were blotted for 6–8 s under 90% relative humidity and plunge-frozen into liquid ethane using a Leica GP2 plunge freezer.

### Cryo-ET data acquisition and subtomogram averaging

Cryo-ET datasets were acquired using a 300-kV Titan Krios microscope (Thermo Fisher Scientific) fitted with a K3 direct electron detector and a GIF energy filter (Gatan). Automated tilt-series acquisition was controlled through SerialEM ^32^. Images were recorded in superresolution mode at a pixel size of 1.06 Å, using an angular increment of 3° over a −48° to +48° range. A cumulative exposure of ∼68 e−/Å² was applied to each tilt series, and images were collected at a defocus of −4.8-5 μm. Raw movie frames were first corrected for beam-induced motion with MotionCor2 ^33^. The corrected images were then imported into IMOD ^34^, where individual projections were combined to generate aligned tilt stacks. Depending on data quality, alignment was carried out either using fiducial markers or with a fiducial-free procedure. CTF parameters were determined with Gctf ^35^, followed by phase correction using ctfphaseflip in IMOD. Two sets of tomographic reconstructions were generated with Tomo3D ^36^: SIRT reconstructions were used to facilitate visualization and identification of Dot/Icm particles, whereas WBP reconstructions served as the input for subsequent subtomogram analysis. Three-dimensional particle alignment, classification, and averaging were carried out with the protomo package ^37^. Dot/Icm particles were first oriented and aligned using 8×binned tomograms. The resulting particle positions and orientations were then transferred back to the unbinned tomograms for subtomogram extraction, followed by refinement using 4×-binned particles. A final round of alignment was performed with 2×-binned subtomograms to recover higher-resolution structural features.

### *In situ* cryo-EM sample preparation, data acquisition and image processing

To enrich secretion-competent Dot/Icm T4SS complexes, bacterial suspensions were incubated with freshly prepared macrophage lysate prior to vitrification ^13^. Treated bacteria were applied to glow-discharged Quantifoil grids and plunge-frozen under identical conditions. This treatment increased the proportion of secretion-competent complexes exhibiting extracellular channel extensions. Cryo-EM movies were collected using a Titan Krios electron microscope (Thermo Fisher Scientific) operating at 300 kV and equipped with a K3 direct electron detector and GIF energy filter (Gatan). Data were acquired in superresolution mode at a pixel size of 0.534 Å. Multishot acquisition was performed using SerialEM ^32^, with acquisition positions targeted to bacterial cell poles located within ice holes. Each movie was recorded over 40 frames with an exposure time of 0.07 s per frame, corresponding to a total electron dose of 70 e^−^/Å^2^. The nominal defocus range was −1.0 to −2.0 μm. Movie alignment and CTF estimation were subsequently carried out in CryoSPARC ^38^. The image-processing workflow is summarized in Extended Data Fig. 5 and Extended Data Table 3).

### Purification of the inner membrane complex (IMC)

To purify DotO and its associated IMC, a C-terminal FLAG tag was introduced into the chromosomal *dotO* locus in a *L. pneumophila* Δ*dotB* mutant background. Briefly, 1,000-bp regions flanking the *dotO* stop codon were cloned into the suicide vector pSR47S to generate a tagging construct in which the FLAG-encoding sequence was inserted immediately before the stop codon. Cells were grown on charcoal yeast extract (CYE) agar plates and subsequently inoculated into AYE medium supplemented with streptomycin (100 μg/ml). Cultures were expanded to a total volume of 4 L and grown at 37 °C for 48 h to an OD₆₀₀ of 3. Cells were harvested by centrifugation, resuspended in lysis buffer containing 50 mM Tris–HCl (pH 7.8), 300 mM NaCl, and 10% (v/v) glycerol, and disrupted using an EmulsiFlex-C3 high-pressure homogenizer (Avestin). Membranes were pelleted by ultracentrifugation at 40,000 rpm (185,677 × *g*) for 2h at 4 °C using a Type 45 Ti rotor (Beckman Coulter). Membrane proteins were solubilized with 1% (w/v) lauryl maltose neopentyl glycol (LMNG; Anatrace) for 1h at 4 °C, and the soluble fraction was applied to anti-FLAG G1 affinity resin (GenScript). The resin was washed with 20 column volumes of wash buffer containing 25 mM Tris–HCl (pH 7.5), 150 mM NaCl, and 0.02% (w/v) LMNG. Proteins were then eluted with wash buffer supplemented with FLAG peptide (200 μg/ml). The eluate was concentrated and further purified by size-exclusion chromatography on a Superose 6 Increase 10/300 GL column (GE Healthcare) equilibrated with wash buffer. Peak fractions were pooled and concentrated for cryo-EM analysis.

### Cryo-EM sample preparation, data acquisition, and image processing of purified IMC

Aliquots (3 μl) of the purified complex at ∼1.5 mg/ml were applied to glow-discharged Quantifoil R1.2/1.3 300-mesh copper grids. Grids were blotted for 3.5 s at 10 °C and 90% relative humidity and plunge-frozen in liquid ethane using a Leica EM GP2 plunge freezer. Cryo-EM data were collected on a Titan Krios electron microscope (Thermo Fisher Scientific) operated at 300 kV and equipped with a K3 direct electron detector and a GIF energy filter (Gatan). Movies were recorded at a nominal magnification of 81,000× in super-resolution mode, corresponding to a pixel size of 0.534 Å. Each movie comprised 40 frames with a total electron exposure of 50 e^-^/Å², with defocus range was -1.0 to -2.0 μm. The image-processing workflow is summarized in Extended Data Fig. 2. Motion correction and contrast transfer function (CTF) estimation were performed in cryoSPARC. Manually picked particles were used to train a Topaz model for automated particle picking. A total of 4,179,701 particles were extracted from 7,234 micrographs. Following 2D classification, 754,579 particles were retained and used for ab initio reconstruction and subsequent heterogeneous refinement with global alignment. Further heterogeneous refinement identified a subset of 339,290 particles, which was subjected to non-uniform (NU) refinement followed by local refinement, yielding a reconstruction at an overall resolution of 3.33 Å. To mitigate the effects of preferred particle orientation, the dataset was further processed using cryoPROS ^39^, resulting in a reconstruction from 135,622 particles at an overall resolution of 3.61 Å. The resulting density map was further processed using EMReady2 ^40^ for density enhancement.

### Model building and refinement

The AlphaFold-predicted structures were used as the initial model for atomic model building. The model of the DotO–DotI–DotJ complex with a stoichiometry of 2:3:1 was also predicted to guide model building. Individual components were fitted into the cryo-EM density map and manually adjusted in Coot v0.8.9.1 ^41^. Residue assignments and side-chain conformations were inspected and corrected according to the density. The model was refined iteratively in real space using Phenix v1.21 ^42^ with secondary-structure and geometry restraints, alternating with manual adjustments in Coot. The final model was validated using Phenix ^42^ and MolProbity ^43^.

### Identification and validation of Porin III

To identify Porin III, a density-derived model from our previously determined *in situ* structure was used as a query for structural homology searches against a library of AlphaFold-predicted *L. pneumophila* protein structures using a locally installed DALI server ^44^. Lpg0731 was identified as the highest-scoring match. To test this assignment, a *lpg0731* transposon insertion mutant was generated and analyzed by *in situ* cryo-EM. A total of 4,853 micrographs were collected from which 5,748 Dot/Icm T4SS particles were identified. Reconstruction of the outer membrane complex with C13 symmetry imposed yielded a map at 3.81 Å resolution. Comparison with the wild-type reconstruction revealed the loss of density corresponding to Porin III, supporting its assignment as Lpg0731.

### Macrophage culture and infection for cryo-FIB milling

THP-1 monocytes were maintained in RPMI 1640 medium supplemented with 10% fetal bovine serum at 37 °C in 5% CO₂. For cryo-FIB/ET, glow-discharged Quantifoil Au R2/2 grids were placed in tissue culture dishes and THP-1 cells were seeded directly onto the grids. Monocytes were differentiated into macrophage-like cells by incubation with 100 ng/mL phorbol 12-myristate 13-acetate (PMA) for 48 h. Differentiated macrophages were infected with *L. pneumophila* LP01 at a multiplicity of infection (MOI) of 20 for 1 h before vitrification.

### Segmentation and 3D Visualization

Tomogram segmentation was performed using Dragonfly 3D World (v. 2022.2), and the resulting segmented structures were rendered and visualized using Blender (v. 5.1.2). Blender was also used to generate three-dimensional animations and supplementary movies.

## Data availability

The atomic coordinates and corresponding density maps have been deposited in the Protein Data Bank (PDB) and the Electron Microscopy Data Bank (EMDB). DotO dimer structure: PDB 39FO and EMD-79402; Overall Dot/Icm T4SS structure: PDB 39GN; Dot/Icm T4SS in the resting state: EMD-79604. Dot/Icm T4SS in the activated state: EMD-79530.

## Acknowledgments

This work was supported by grant R01AI152421 (J.L. and C.R.R.) from National Institute of Allergy and Infectious Diseases (NIAID) and National Institutes of Health (NIH). S.S., H.W., and W. M. were supported by grant R01GM137068 (W.M.) from National Institute of General Medical Sciences (NIGMS) and NIH. Cryo-EM data were collected at the Yale CryoEM Resource funded in part by the NIH grant 1S10OD023603-01A1. We thank Jennifer Aronson for critical reading of the manuscript and helpful comments.

## Supplemental Information

**Extended Data Fig. 1.**
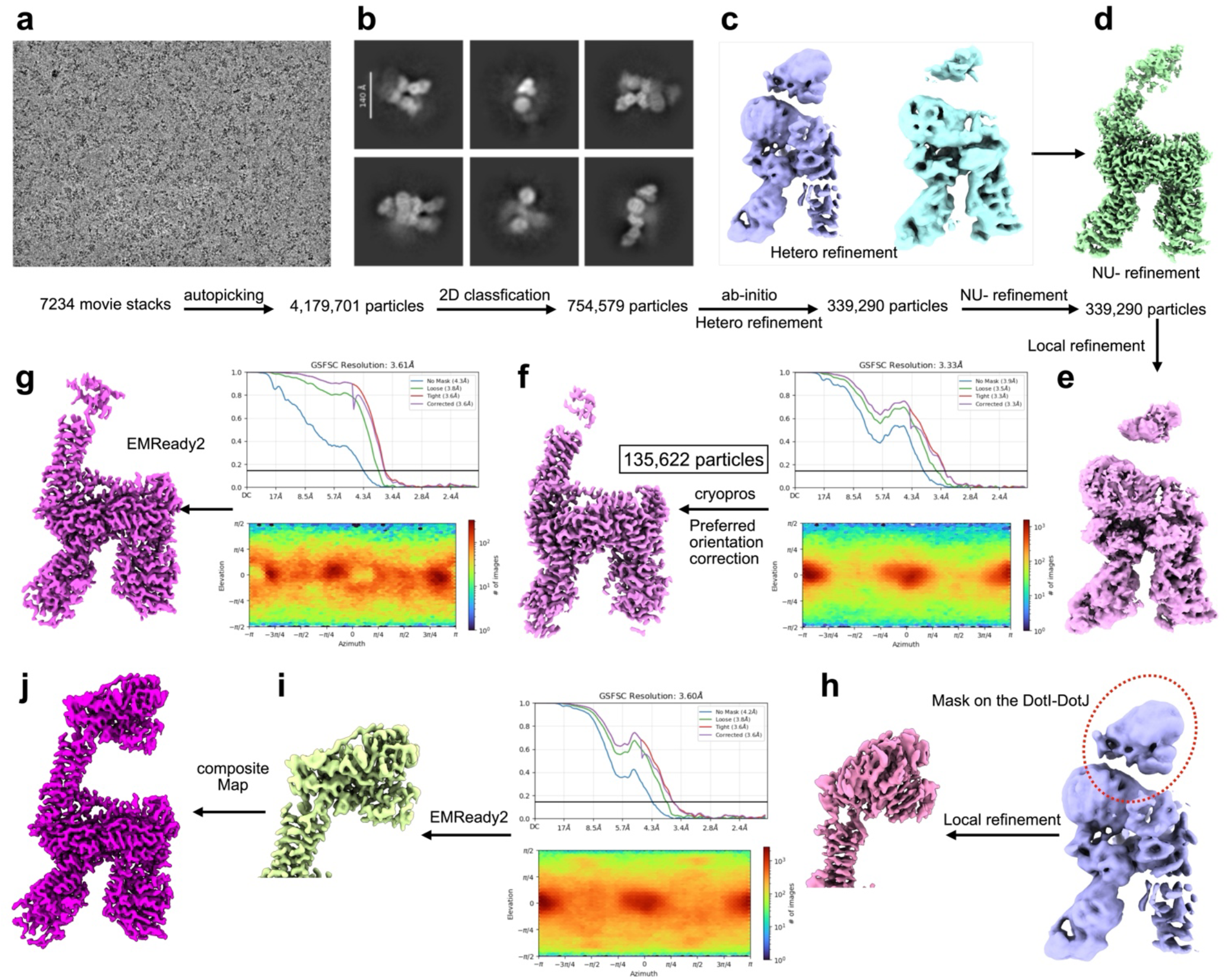
Cryo-EM data-processing workflow for the IMC complex. **a**, Representative cryo-EM micrograph from 7,234 movie stacks. **b**, Two-dimensional (2D) class averages. Automated particle picking yielded 4,179,701 particles, of which 754,579 were retained after 2D classification. **c**, Heterogeneous refinement identified a subset of 339,290 particles for further refinement. Representative heterogeneous-refinement maps are shown. **d**, Reconstruction obtained by non-uniform (NU) refinement. **e**, Subsequent local refinement yielded a reconstruction at a nominal resolution of 3.33 Å. **f**, Preferred-orientation correction using cryoPROS yielded a reconstruction from 135,622 particles at a nominal resolution of 3.61 Å. Gold-standard Fourier shell correlation (FSC) curves and particle angular distributions are shown for the reconstructions before and after correction. **g**, Map obtained following EMReady2 enhancement of the orientation-corrected reconstruction. **h**, focused local refinement of the DotI–DotJ region, indicated by the red dashed outline, yielded a reconstruction at 3.60 Å; the corresponding FSC curves and particle angular distribution are shown. **i**, EMReady2-enhanced map of the locally refined DotI–DotJ region. **j**, Final composite map assembled from the maps shown in **g** and **i**. Nominal resolutions were estimated using the gold-standard FSC threshold of 0.143.

**Extended Data Fig. 2.**
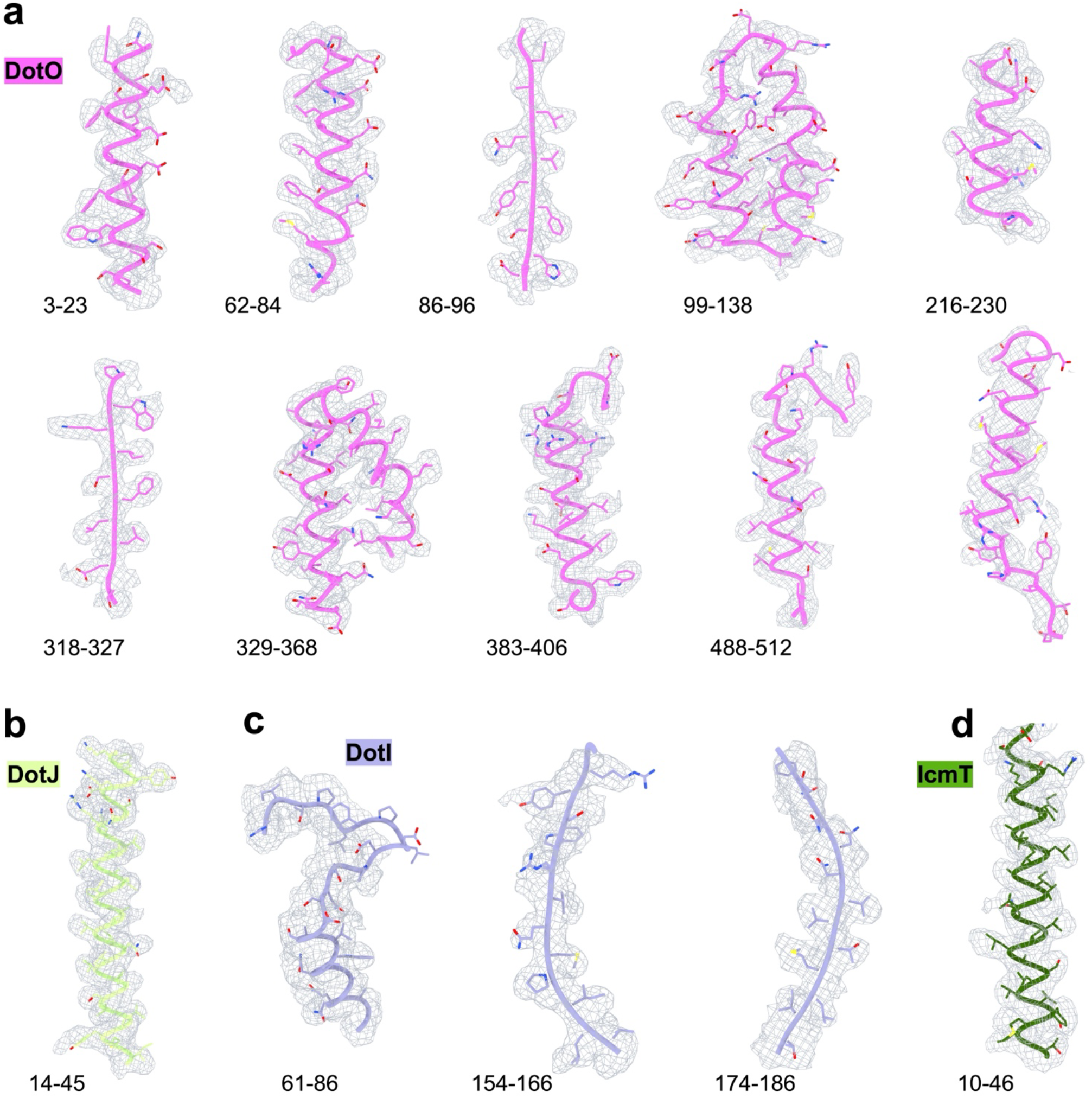
Representative cryo-EM densities and models fitting. **a–d**, Selected regions of the models shown within the corresponding cryo-EM density (gray mesh). Models are displayed as cartoons and sticks. Residue ranges are indicated below each view.

**Extended Data Fig. 3.**
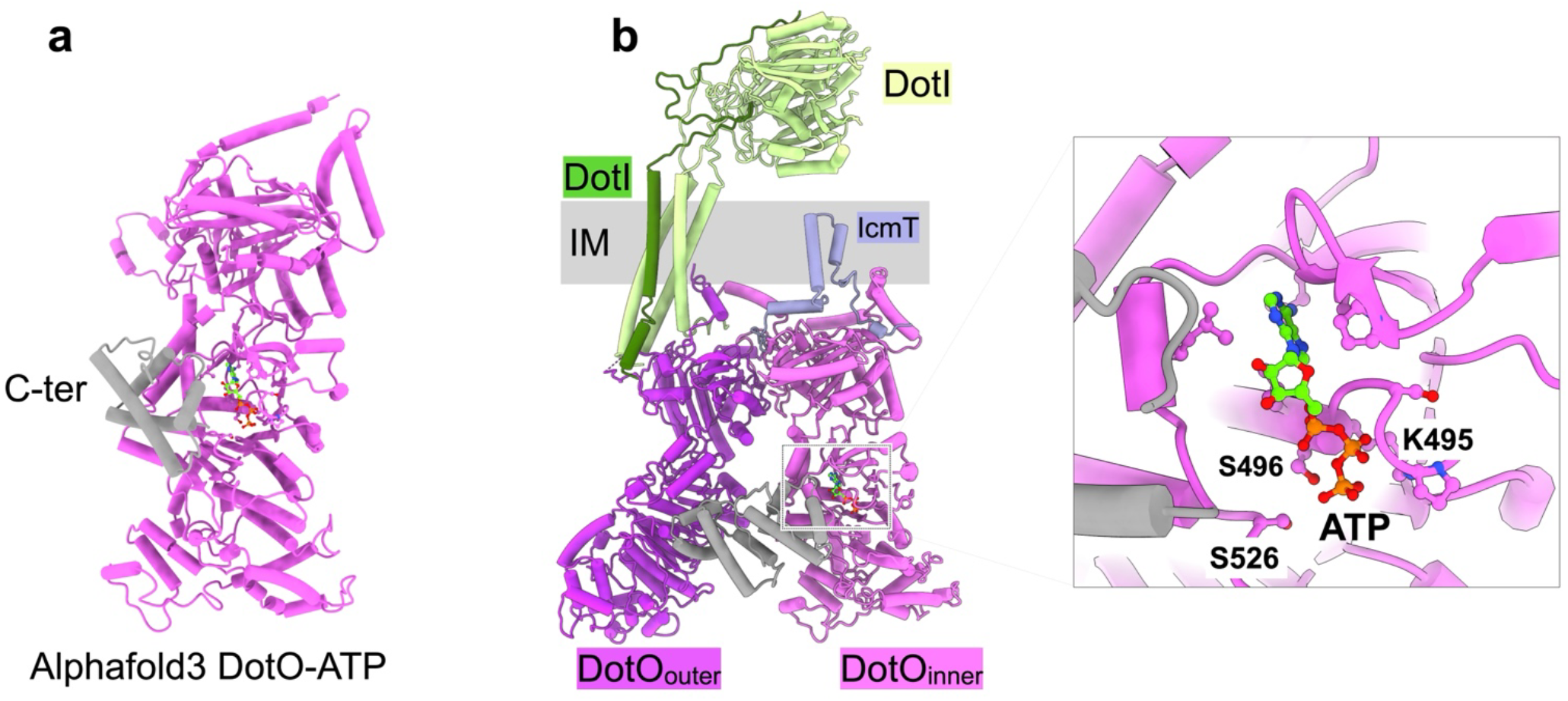
Predicted ATP-binding site of DotO. **a**, AlphaFold3 model of ATP-bound DotO, with the C-terminal region shown in gray. **b**, Location of the predicted ATP-binding site within the IMC complex. The enlarged view shows ATP and nearby residues K495, S496, and S526. IM, inner membrane.

**Extended Data Fig. 4.**
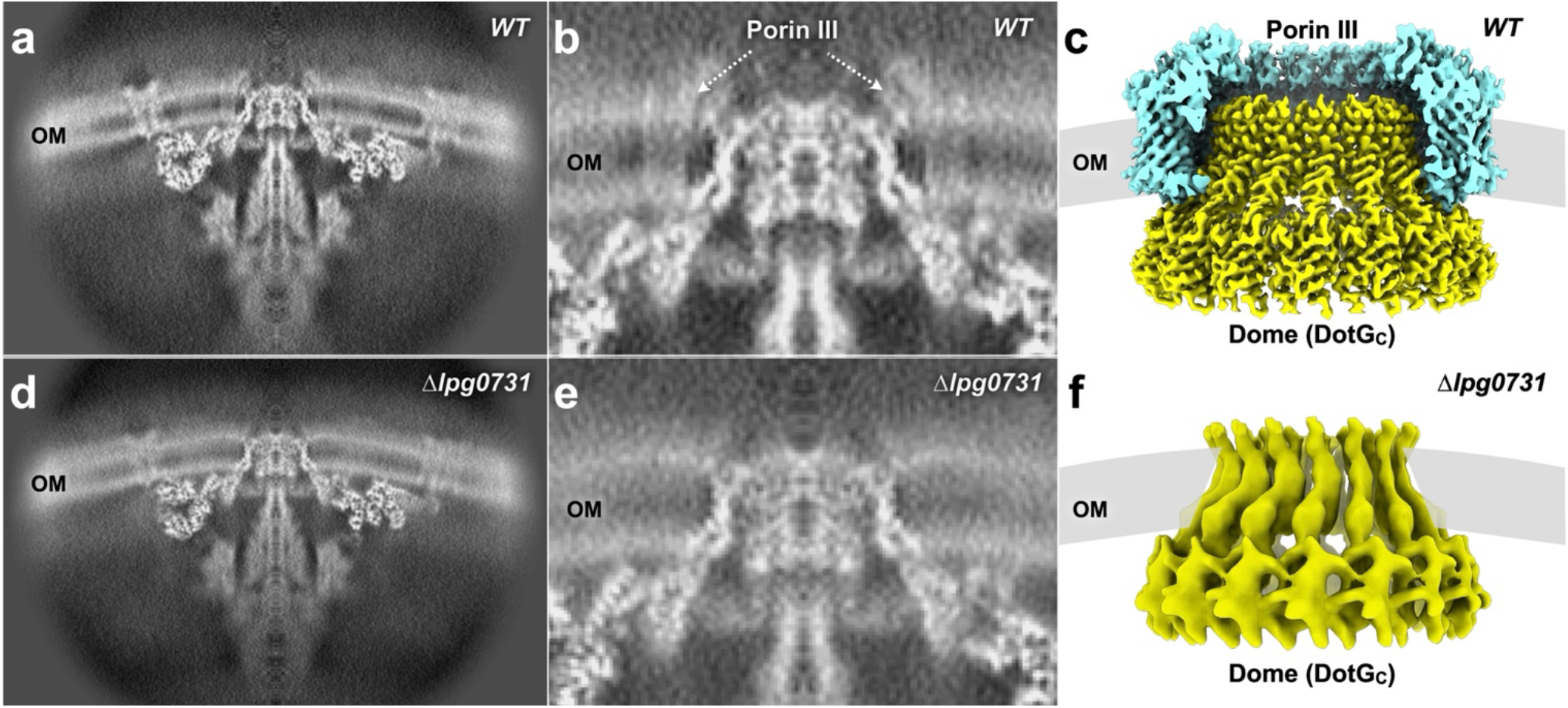
Loss of *lpg0731* eliminates the Porin III density surrounding the DotG dome. **a**,**d,** Central sections through *in situ* cryo-EM reconstructions of the Dot/Icm T4SS from wild-type (WT; a) and Δ*lpg0731* (d) cells. **b**,**e,** Enlarged views of the outer-membrane region showing the Porin III density in WT (b) and its absence in Δ*lpg0731* (e). **c**,**f,** Surface rendering representations of the outer-membrane region in the *dotB* E191K (c) and Δ*lpg0731* (f) complexes, showing Porin III (cyan) surrounding the DotG C-terminal dome (yellow) in the former and the retained dome without the surrounding ring in the latter. These observations identify Lpg0731 as a component of the Porin III ring. OM, outer membrane.

**Extended Data Fig. 5.**
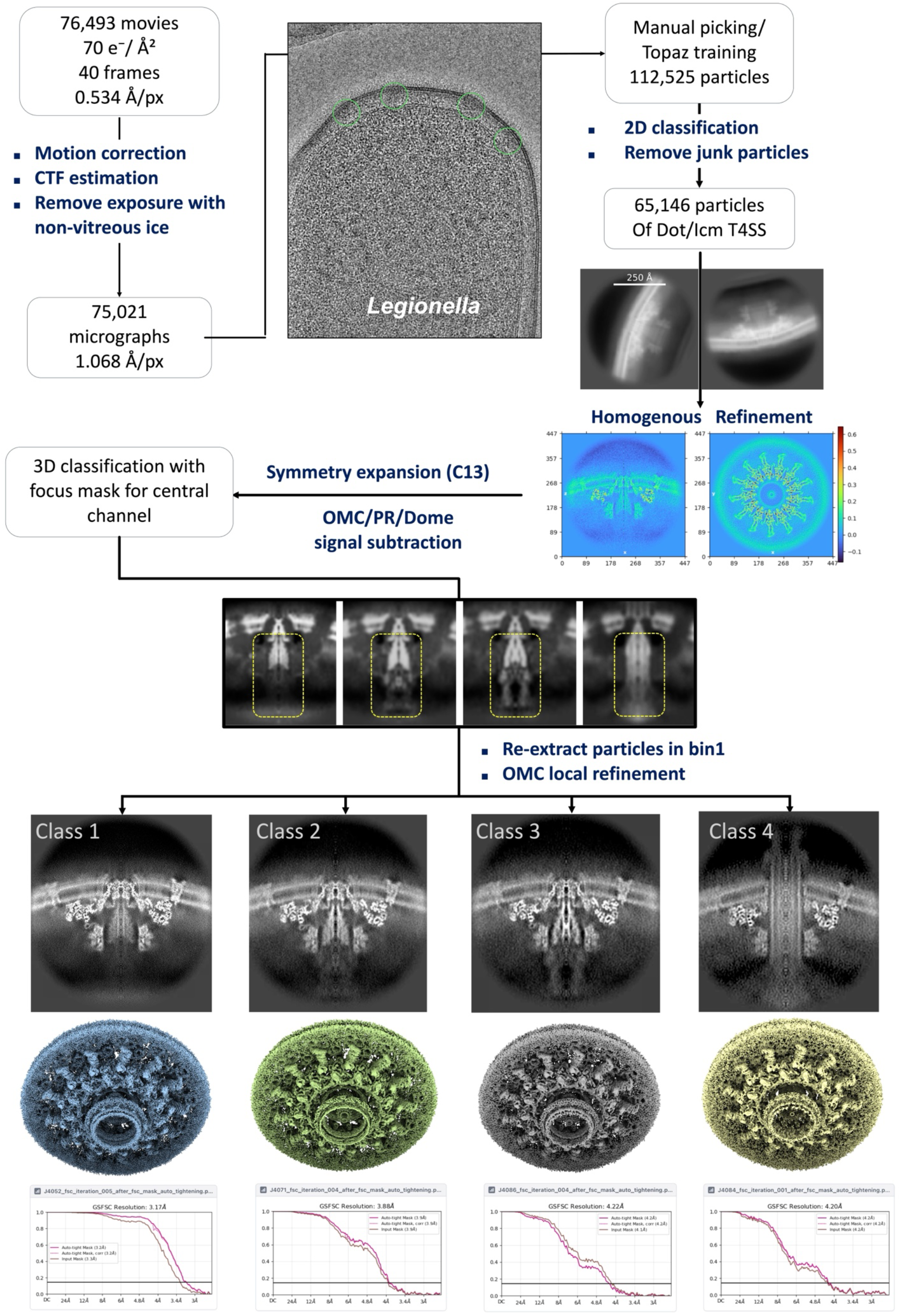
Cryo-EM data processing workflow for the Dot/Icm T4SS from wild-type *L. pneumophila* following incubation with host-cell lysate. A total of 76,493 movies were collected and subjected to motion correction and contrast transfer function (CTF) estimation. Micrographs containing non-vitreous ice were excluded, yielding 75,021 micrographs for further processing. The central panel shows a representative cryo-EM micrograph of wild-type *L. pneumophila*, with identified particles highlighted by green circles. Manual particle picking and Topaz-based particle selection yielded 112,525 particles. Following twofold-binned (bin 2) particle extraction and 2D classification to remove low-quality particles, 65,146 Dot/Icm T4SS particles were retained. Initial homogeneous refinement yielded a reconstruction at 3.78 Å resolution. The OMCC, PR and dome densities were subsequently subtracted, followed by C13 symmetry expansion and focused 3D classification of the central secretion channel. After removal of duplicate particles, the remaining particles were re-extracted at full resolution (bin 1) and subjected to local refinement of the OMCC and CTF refinement. The resulting four structural classes are shown with their corresponding cryo-EM reconstructions and Fourier shell correlation (FSC) curves, yielding resolutions of 3.17 Å (class 1), 3.88 Å (class 2), 4.22 Å (class 3) and 4.20 Å (class 4).

**Extended Data Fig. 6.**
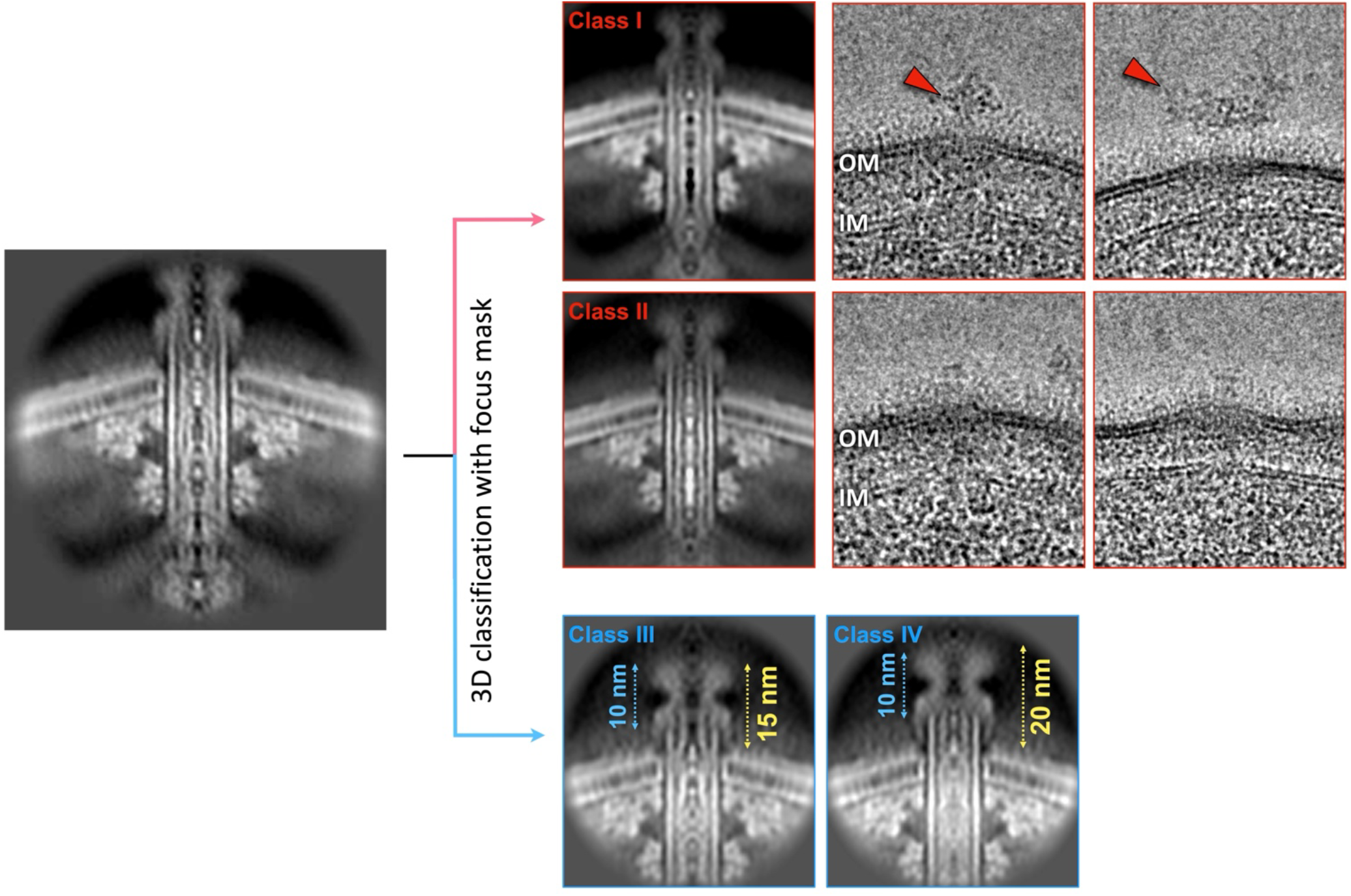
Focused classification reveals distinct conformations of the active Dot/Icm T4SS secretion channel. The overview (left) highlights the region selected for focused classification. Classes I and II (red) represent empty and putatively substrate-occupied secretion channels, respectively. For each class, the left panel shows the classified T4SS reconstruction, whereas the panels on the right show representative particles mapped back to their original cryo-EM micrographs. In Class I (top), the secretion channel is empty, with additional density above the apparatus (red arrowheads) potentially representing substrate released from the channel. In Class II (middle), additional density within the channel is consistent with a translocating substrate. Classes III and IV (blue; bottom) reveal variations in the length of the extracellular channel extension. Arrows indicate the measured dimensions, including a common 10-nm region and extracellular extensions of 15 nm in Class III and 20 nm in Class IV. OM, outer membrane; IM, inner membrane.

**Extended Data Table 1.** Cryo-ET data collection parameters.

| Data collection parameter |  |
| --- | --- |
| Microscope | Titan Krios (Thermo Fisher Scientific) |
| Accelerating voltage (kV) | 300 |
| Nominal magnification | 42,000× |
| Physical pixel size (Å) | 2.148 |
| Total electron dose (e<sup>-</sup>/Å<sup>2</sup>) | 68 |
| Frame Number | 10 |
| Acquisition scheme | -48 /+48, 3° |
| Defocus range (µm) | -4.8 to -5.3 |
| Detector | Gatan K3 |
| Energy-filter slit width (eV) | 20 |
| Data processing |  |
| Total tomograms | 403 |
| Software packages | IMOD, Etomo, Tomo3D, I3 |
| Total T4SS particles | 1499 |
| Number of subtomograms after classification | 970 (resting state)- 529 (529) |
| Symmetry imposed | C13 |
| Resolution | ~25 Å |

**Extended Data Table 2.** CryoEM data collection and structure refinement statistics.

|  | DotO Dimer | Overall Dot/Icm T4SS structure |
| --- | --- | --- |
| Data collection |  |  |
| Microscope | Titan Krios |  |
| Voltage (kV) | 300 |  |
| Camera | K3 |  |
| Magnification | 81K |  |
| Total Electron Exposure (e<sup>-</sup>/Å<sup>2</sup>) | 50 |  |
| Defocus range (um) | -1.2 to -2.2 |  |
| Pixel size (Å) | 1.068 |  |
| Micrographs (no.) | 7,234 |  |
| Data processing |  |  |
| Symmetry imposed | C1 |  |
| Final particle images (No.) | 135,622 |  |
| Map resolution (Å) 0.143 FSC threshold | 3.61 |  |
| Map sharpening B factor (Å<sup>2</sup>) | 101.7 |  |
| Refinement |  |  |
| CC (model vs. data) | 0.80 | / |
| Non-hydrogen atoms | 21,664 | 576,720 |
| Protein residues | 2,768 | 73,617 |
| Ligands | 0 | 0 |
| B factors (Å<sup>2</sup>) |  |  |
| Proteins (min/max/mean) | 97.59 | / |
| R.m.s. deviations |  |  |
| Bond lengths (Å) | 0.003 | 0.004 |
| Bond angles (°) | 0.660 | 0.86 |
| Validation |  |  |
| MolProbity score | 2.07 | 1.94 |
| Clashscore | 4.60 | 4.11 |
| Rotamer outliers (%) | 4.46 | 3.85 |
| Ramachandran plot |  |  |
| Favoured (%) | 95.09 | 94.53 |
| Allowed (%) | 4.80 | 5.30 |
| Outliers (%) | 0.11 | 0.18 |
| PDB code | 39FO | 39GN |
| EMDB code | EMD-79402 |  |

**Extended Data Table 3.** *In-situ* cryoEM data collection and structure refinement statistics.

|  | Overall | Resting<br>state | Pre-primed<br>state | Primed<br>state | Active<br>state |
| --- | --- | --- | --- | --- | --- |
| Data collection |  |  |  |  |  |
| Microscope | Titan Krios | Titan Krios | Titan Krios | Titan Krios | Titan Krios |
| Voltage (kV) | 300 | 300 | 300 | 300 | 300 |
| Camera | K3 | K3 | K3 | K3 | K3 |
| Magnification | 81K | 81K | 81K | 81K | 81K |
| Total Electron Exposure (e<sup>-</sup>/Å<sup>2</sup>) | 70 | 70 | 70 | 70 | 70 |
| Defocus range (um) | -1.2 to -2.2 | -1.2 to -2.2 | -1.2 to -2.2 | -1.2 to -2.2 | -1.2 to -2.2 |
| Pixel size (Å) | 1.068 | 1.068 | 1.068 | 1.068 | 1.068 |
| Total micrographs |  | 76,493 |  |  |  |
| Data processing |  |  |  |  |  |
| Reconstruction | Global | Local | Local | Local | Local |
| Final particle images | 65,146 | 43,902 | 11,239 | 5,640 | 4,365 |
| Box size (Å) | 560 | 560 | 560 | 560 | 560 |
| Fourier size (Å) | 448 | 448 | 448 | 448 | 448 |
| Symmetry imposed for local refinement | C13 | C13 | C13 | C13 | C13 |
| Map resolution OMCC (Å) | 3.78 | 3.17 | 3.88 | 4.22 | 4.20 |
| EMDB code |  | EMD-79604 |  |  | EMD-79530 |

**Movie S1. Animation of the primed state of the Dot/Icm T4SS.**

**Movie S2. Animation of the active state of the Dot/Icm T4SS.**

**Movie S3. Animation of the Dot/Icm mediated bacterium–host interaction.**

